# Juveniles exhibit lower heatwave tolerance than adults in a reef-building coral

**DOI:** 10.64898/2026.09.23.753667

**Authors:** Cinzia Alessi, Liam Lachs, Adriana Humanes, Helios Martinez, Nicolas Croft, Leevan Manuel, Peter J. Mumby, Adeeshia I. Tellei, Daniel Cassidy, James Guest

## Abstract

Marine heatwaves cause widespread coral bleaching and mortality, yet thermal responses often differ between adult and juvenile corals, with the latter frequently being more tolerant. Whether the observed enhanced tolerance of juveniles reflects intrinsic physiological differences across life stages or simply reduced environmental stress due to shaded microhabitats occupied by juveniles, or a combination of both factors, remains unclear. To test whether colony size determines thermal responses and if a threshold size exists for tolerance, we exposed 89 colonies of the reef-building coral *Acropora* aff. *digitifera* to a 6-week, 32.5 °C simulated marine heatwave under controlled light conditions, with cumulative stress reaching 20 °C-weeks. Contrary to expectations, ∼2 °C-weeks less heat stress was required to trigger bleaching and mortality in coral juveniles compared to adults. Mortality divergence across colony sizes was greatest at ∼10 °C-weeks, when the smallest corals exceeded 80% mortality, whereas the largest colonies showed <10% mortality. A breakpoint in survival odds was apparent at 13.4 cm diameter, corresponding to the size at onset of sexual maturity in the native wild population for this species. These results indicate an ontogenetic shift in physiological heat tolerance, with larger reproductively mature colonies showing enhanced tolerance. The apparent survival advantage of juveniles in the reef frequently reported during bleaching events could be more likely due to micro-environmental buffering than intrinsic physiological tolerance. Our findings suggest that juvenile corals may be more vulnerable to climate change than previously thought, particularly as declining habitat complexity could reduce microhabitat refugia on future reefs.

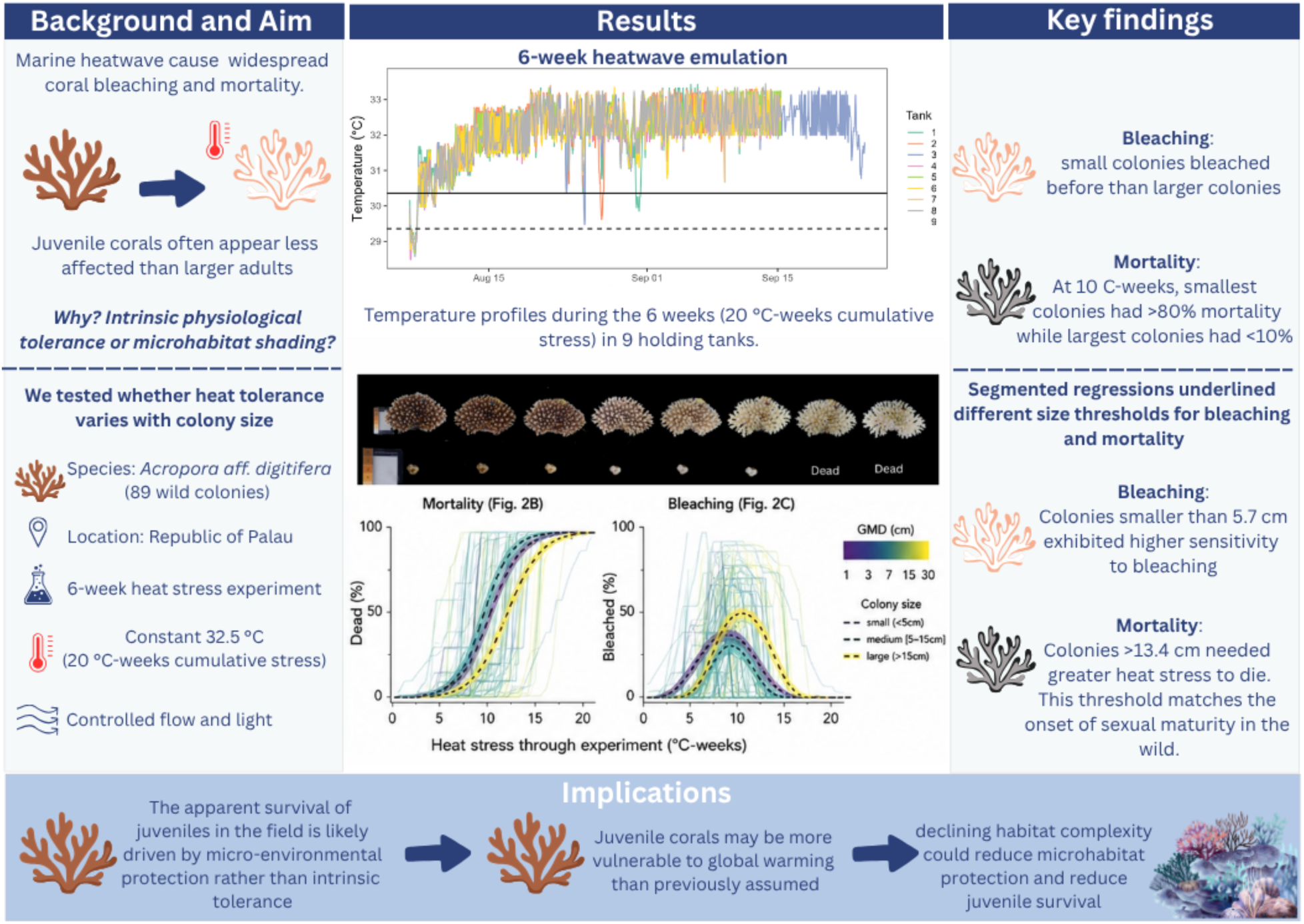

## Introduction

Colony size is critical to Scleractinia coral fitness, directly influencing their growth, competitive ability, survival, and reproduction. Consequently, disturbances that selectively impact specific size classes can fundamentally alter the trajectory and demographic resilience of coral populations.

Ocean warming is presently the principal driver of coral mortality at a global scale, with marine heatwaves altering population size structures (Ceccarelli et al., 2026; Hughes et al., 2003, 2017, 2018; Reimer et al., 2024). Field surveys conducted during and after bleaching events consistently report lower bleaching prevalence among juvenile colonies relative to conspecific adults, suggesting the existence of size-dependent thresholds in thermal sensitivity (Álvarez-Noriega et al., 2018; Burn et al., 2023; Mumby, 1999). However, it remains unclear whether these patterns are driven by size-dependent physiological differences, environmental buffering associated with the microhabitat refugia occupied by small colonies, or a combination of both. Distinguishing between these mechanisms is increasingly important, as climate change is reducing reef structural complexity (Alvarez-Filip et al., 2009), potentially diminishing natural refugia for juveniles and increasing their exposure to stress.

Existing population models assume that juveniles escape bleaching mortality during heatwaves (Bozec et al., 2025; Cresswell et al., 2024; Lachs, 2025; Lachs, Bozec, et al., 2024) in line with field observations (Bena & Van Woesik, 2004; Burn et al., 2023; Mumby, 1999). However, the mechanistic basis of such juvenile heatwave survival is unknown. If higher heat tolerance of juveniles has a purely physiological basis, then we could expect this tolerance in juveniles to persist in the future. However, if juveniles survive only due to environmental buffering from their microhabitat refugia, then their low levels of heatwave mortality will depend on reef structural complexity. If reef structure is lost in the future due to reef degradation, juveniles could be increasingly exposed to heatwave impacts. Therefore, resolving the mechanistic basis of size-dependent bleaching responses will help to improve ecological forecasts under climate change.

Juvenile corals initially grow as small, encrusting colonies that spread over the reef substrate (Ritson-Williams et al., 2009), a morphology thought to enhance mass-transfer efficiency and thereby potentially increase thermal tolerance (van Woesik, Irikawa, et al., 2012). In adult colonies, morphological plasticity interacts with physiological plasticity to actively optimize photosynthetic energy acquisition rather than simply maximizing light capture (Hoogenboom et al., 2008), raising the possibility that juvenile morphology similarly shapes energy acquisition efficiency alongside gas exchange. Any such benefit, however, is constrained by colony size; Energetic models predict that net energy surplus scales positively with colony size as energy intake scales linearly with surface area. Consequently, small juveniles typically operate with a smaller absolute energy surplus than larger, mature colonies despite the substantial reproductive investments of larger mature individuals (Leuzinger et al., 2003; Sebens, 1987). Juveniles may therefore face a trade-off between two competing effects, as their limited surface area may constrain total energy acquisition, so any thermal-tolerance benefit of this early allocation window likely depends on how efficiently their smaller energy budget is directed toward metabolic maintenance and growth, rather than on its absolute size (van der Meer, 2006).

Testing how heat tolerance varies with colony size in the field is inherently challenging because ontogenetic differences in physiology are confounded by differences in microhabitat where colonies establish. Coral larvae preferentially settle in cryptic, shaded microhabitats where irradiance and hydrodynamic conditions differ markedly from those experienced by exposed adult colonies on the reef (Doropoulos et al., 2015; Mumby, 1999). These microhabitat differences can have direct implications for how corals physiologically respond to their environment. Elevated irradiance and thermal stress act synergistically to promote photoinhibition and the accumulation of reactive oxygen species (ROS) in coral tissue, while water flow directly affects the thickness of the boundary layer and coral mass transfer (Brown, 1997; Comeau et al., 2019; Nakamura et al., 2005). Consequently, juveniles and adults often experience markedly different thermal and light environments, making it difficult to determine whether observed differences in thermal performance reflect intrinsic physiological properties or environmental variation.

To disentangle these effects, we address this knowledge gap by exposing whole colonies of *Acropora* aff. *digitifera* spanning a broad size range to a controlled marine heatwave emulation under standardized microhabitat conditions of irradiance and flow. This study aimed to quantify the influence of coral colony size on heatwave-driven bleaching and mortality, whilst controlling habitat variability that can confound field observations of size-dependent heatwave impacts. These insights are important for understanding how early life stages respond to heatwaves and their ecological and evolutionary implications under climate change.

## 2 Material and Methods

### 2.1 Coral collection

The reef-building *Acropora* aff. *digitifera* was chosen as a model species for this study due to its abundance in the shallow water reefs of the Indo-Pacific region. *A.* aff. *digitifera* is a morphologically distinct corymbose species, for which extensive heat tolerance and demographic research has been conducted in the Republic of Palau (Humanes et al., 2021, 2024; Lachs et al., 2023; van der Steeg et al., 2025). In July 2025, 104 colonies ranging in size (Fig. 1B, 1C) were collected using a stratified design targeting ∼10 individuals per size class across five categories (0-5 cm, 5-10 cm, 10-15 cm, 15-20 cm, 20-25 cm, and 25-30 cm). Colonies were collected from the Mascherchur Reef (GPS: 7.285632, 134.527444, Palau, Fig. 1A, C) using a hammer and chisel and brought to the Palau International Coral Reef Center (PICRC), where the experiment was carried out. The distance between the chosen colonies was at least 5 m to maximize the chance of sampling distinct genets rather than clones.

**Fig. 1.**
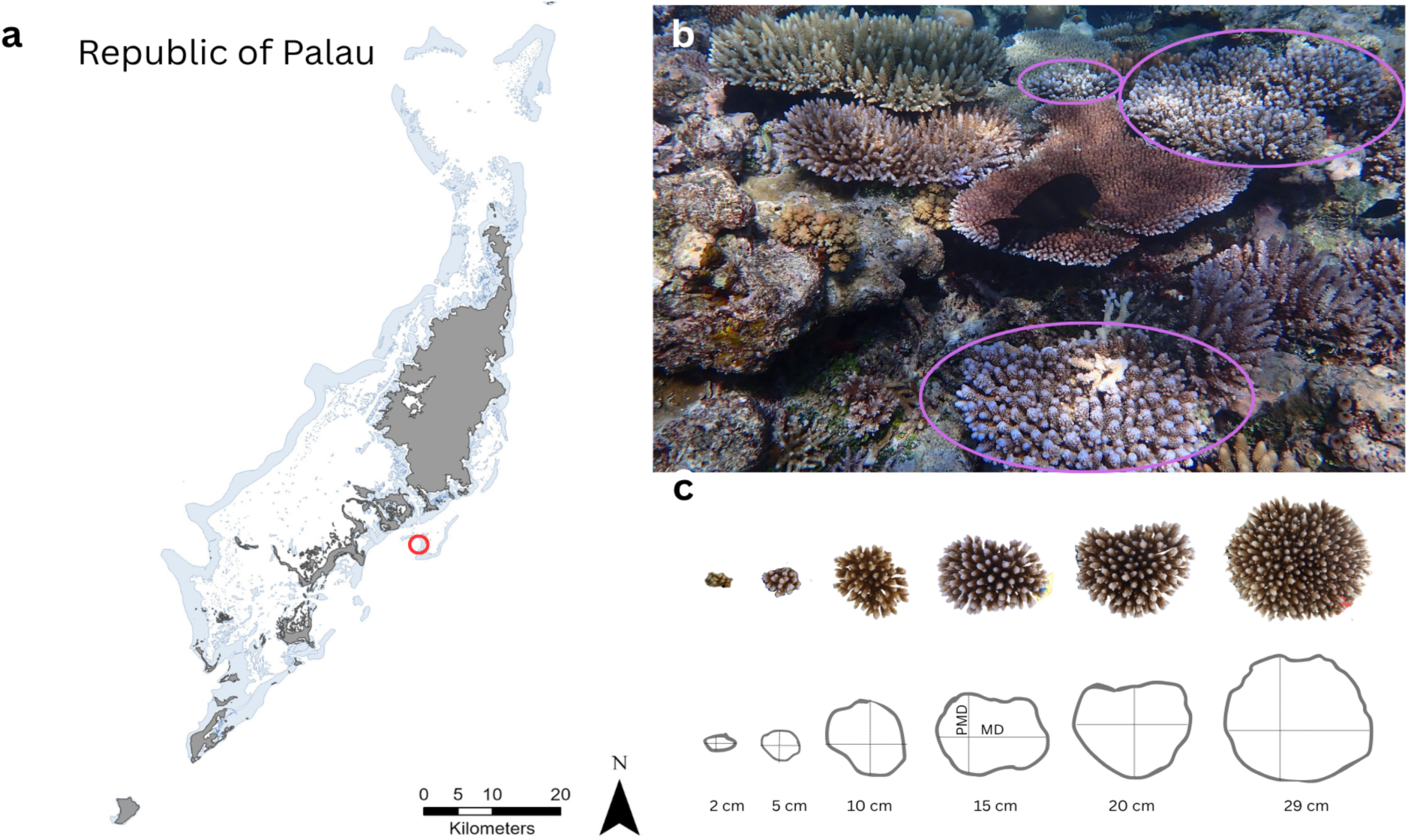
Map of the Republic of Palau showing the collection site of *Acropora* aff*. digitifera* colonies at Mascherchur Reef (Red circle, a). Representative image of the reef habitat at Mascherchur Reef, with purple circles indicating wild *A.* aff*. digitifera* colonies (b). Representative colony morphologies across size classes, illustrating the measurements used to calculate Geometric Mean Diameter (GMD), including Maximum Diameter (MD) and Perpendicular Maximum Diameter (PMD) (c).

Once at PICRC, colonies were placed in an outdoor holding tank (760 L), spaced evenly to allow flow between colonies, and positioned on individual stands. Water oxygenation was maintained with a constant water inflow, and three pumps (EHEIM Universal 1200, EHEIM GmbH & Co. KG, Germany) ensured continuous water circulation. Colonies were kept under these conditions for 12 days to allow acclimatization and recovery after collection. During this period, each colony was photographed, measured, and the main taxonomic features such as axial and radial corallites were re-inspected to confirm species morphological characteristics, and cataloged with a unique ID. After this verification, 15 colonies were removed due to different morphological features than *A.* aff. *digitifera*. The maximum and perpendicular diameters were measured for each colony to calculate the geometric mean diameter (GMD; Guest et al., 2014).

### 2.2 Experimental design and normalization of environmental variables

In August 2025, following colony collection, 89 colonies of various sizes were distributed across nine experimental tanks (77L x 52W x 36H cm, 102L), ensuring that each tank contained a comparable range of colony sizes. Stands of varying heights were made from egg crates to ensure the smaller colonies received equivalent light and water flow as adult colonies. Each tank was equipped with two pumps (Rio+ 400, 10W; TAAM Inc., Taiwan) to ensure water circulation in each tank and three lights (48” 50/50 XHO LED, Reef Brite Ltd., USA) under a 12:12 h light-dark cycle. Light intensity was measured at six different points in each tank using a LI–250A light meter (Licor), and it was adjusted to be 181 ±µmol photons m^-2^ s^-1^ at the bottom of the tank.

Temperature was regulated through a constant inflow of both ambient temperature and warm water. The ambient temperature sump was directly connected to the aquarium’s main pipeline and supplied all tanks, whereas each tank had its own sump, equipped with two pumps and two heaters controlled by two independent controllers. This system allowed precise temperature control throughout the long-term marine heatwave emulation experiment following Humanes et al. (2024). To measure temperatures, Hobo pendant data loggers with a recording time set at a 10-minute interval were placed in each tank. The accumulation of heat stress was measured in terms of Degree Heating Weeks (DHW), a metric developed by the National Oceanic and Atmospheric Administration’s (NOAA) Coral Reef Watch (CRW) which captures both the intensity and duration of heat exposure (Skirving et al., 2020). To quantify DHWs in our experiment, we used a locally derived in situ-adjusted MMM for the source reef derived from in situ and satellite data comparisons (Lachs, Humanes, et al., 2024) to then compute experimental DHWs following Humanes et al. (2024).

Water temperature increased in each tank following a ramp profile. The temperature rose by 0.5°C every two days from 30.5°C on day 1 (August 7^th^) to a maximum of 32.5°C on day 13 (August 19^th^), where temperatures were then maintained until the end of the experiment when all the individuals reached mortality (September 24^th^). The in situ adjusted Maximum Monthly Mean (MMM) baseline temperature in Mascherchur Reef was 29.36°C, placing the bleaching threshold (MMM+1°C) at 30.36°C, which was exceeded from the first day of the ramp. Mean water temperatures were consistent across all tanks throughout the experiment, ranging from 32.07 to 32.15°C overall, with mean daily maxima between 32.70 and 32.85°C and mean daily minima between 31.25 and 31.45°C (Table S1, Fig. S1).

### 2.3 Bleaching-mortality heatwave tolerance

Bleaching and mortality heatwave tolerance was measured following methods described in previous work for the same species (Humanes et al., 2024; Lachs et al., 2024; Humanes et al., 2022; Lachs et al., 2023) but with adjustments made for the fact that our study was conducted on whole colonies rather than colony fragments. Each day of the heat stress experiment, coral colonies were scored as healthy, pale, bleached, or dead, and the percentage area of the colony that fell into each category was recorded. Given that loss of microalgal symbionts can progress extensively before any paling is even visible to the naked eye (Humanes et al., 2022), we considered pale fragments as healthy. These health status categories are strongly predictive of chlorophyll and carotenoid concentrations and the tissue population density of dinoflagellate algal symbionts, fully described in previous work (Humanes et al., 2022). The proportion of colonies in each health state was calculated relative to the total number of colonies and plotted as stacked bar charts across days since the start of the heat stress experiment, both pooled across all colonies and stratified by size class (small: <5 cm, medium: 5–15 cm, large: >15 cm GMD). To explore relationships between cumulative heat stress and colony-level bleaching and mortality, individual colony trajectories were plotted against degree heating weeks (DHW), with colony size represented by a continuous scale (GMD in cm).

### 2.4 Size-dependent survival trajectories

To examine the effects of heat stress and colony size on coral mortality, we fitted a generalized linear mixed model (GLMM) with a binomial distribution using the glmmTMB package in R. The probability of mortality was modeled as a function of degree heating weeks (DHW) and geometric mean diameter (GMD), including their interaction term, to assess whether the effect of heat stress on mortality varied with colony size. Colony identity and tank were included as random effects to account for repeated measurements within colonies and potential tank-level variation. Results were visualized as mortality probability curves across the DHW gradient at multiple colony sizes, as line plots of predicted mortality across the GMD gradient at discrete DHW levels (7–12 °C-weeks).

### 2.5 Colony size thresholds for bleaching and mortality heat tolerance

For each coral colony, thermal tolerance was characterized using two endpoints derived from cumulative thermal exposure expressed as degree heating weeks (DHW, °C-wks). The lethal dose of heat stress required to elicit 50% mortality for each colony (LD50) was calculated as the minimum DHW at which cumulative mortality reached or exceeded 50%. The bleaching mortality index ED50 (BMI ED50) was defined as the effective DHW dosage required to cause BMI to reach or exceed 0.5, representing the point at which either half the colony was dead and half alive, or the entire colony was fully bleached. Additionally, the bleaching ED25 and ED50 of each colony were computed as the minimum DHW at which the proportion of bleached tissue reached 25% or 50%, respectively. However, these could not be computed for colonies that never reached the relevant bleaching threshold (e.g., a colony that transitioned directly from healthy to dead).

To examine the relationship between coral colony size and heatwave tolerance, we fitted segmented regression models relating LD50 and ED50 to colony geometric mean diameter (GMD). For each response variable, we first fitted a linear model including GMD and tank as covariates to account for tank effects, then applied segmented regression using the segmented R package to identify a single breakpoint in the relationship with GMD. Fitted values and 95% confidence intervals were extracted using the broken.line() function. The same approach was applied to bleaching ED25 and BMI ED50 endpoints.

### 2.6. Water flow influences colony responses

To evaluate how water flow affected each colony size class, we used the gypsum dissolution method (Jokiel & Morrissey, 1993). Gypsum clod cards of the same shape but different size classes were created to represent small, medium, and large coral colonies. These were placed across the tanks, recreating the spatial distribution of coral colonies in the tanks during the heat stress experiment. After 24 hours, clod cards were sun-dried until a constant weight was obtained. The proportion of weight loss (dissolution rate) was calculated from the difference between initial and final weights. Rates were normalized to the initial weight for comparison among the clod cards of different sizes. The dissolution rate of clod cards was tested using a generalized linear mixed-effects model (GLMM, function glmmTMB) in R, to test whether clod cards of different sizes were exposed to equivalent flow conditions during the experiment. In the model, clod card size was used as a fixed effect, and tank was included as a random intercept. Estimated marginal means were computed using emmeans, and pairwise comparisons among size classes were conducted to assess significant differences.

### 2.7 Additional data

To understand size-dependent reproductive traits, we conducted field reproductive monitoring. We measured colony size and gravidity one week before the full moon in March 2025, preceding the expected *Acropora* mass spawning period in Palau (Gouezo et al., 2020; Penland et al., 2004). A total of eighty-seven colonies were randomly surveyed at Mascherchur Reef (7.285632° N, 134.527444° E; Palau). For each colony, the maximum diameter and the maximum perpendicular diameter were measured to calculate the geometric mean diameter (GMD). The presence or absence of oocytes was determined by cracking two branches in the middle of the colonies to allow visual inspection with a magnifying lens (following Baird et al., 2012). Colonies containing pigmented oocytes were classified as sexually mature. These data were used to fit a logistic regression model with sexual maturity status (binary: 0 = immature, 1 = mature) as the response variable and colony GMD (cm) as the predictor, using a binomial error family (glm, R). This approach allowed the estimation of the colony size at which 50% of colonies are predicted to be sexually mature.

## 3 Results

### 3.1 Environmental variables

The experiment was finished once there were no more surviving corals. The experiment ran for 41 days in all tanks except for tank 3, which ran for 46 days. The maximum DHW accumulated by the end of the experiment was 19.4°C-weeks in tank 3, while the other tanks were turned off 9 days prior at a stress of 15.9 °C-weeks (Table S1).

The dissolution rate of clod cards (Supplementary Fig. S2), as a proxy of likely mass transfer rates of coral colonies, was significantly influenced by size class. Pairwise comparisons confirmed that small-size class clod cards had significantly higher dissolution than medium clod cards (β = 0.21, z = 7.51, *p* = 0.036), whereas differences between small and large (β = -0.06, z = -2.57, *p* = ns) and between medium and large clod cards (β = -0.04, z = -1.6, *p* = ns) were not significant.

### 3.2 Bleaching and mortality response across the experiment

Coral phenotypic status changed progressively with increasing heat stress exposure, with colonies transitioning from healthy through pale and bleached states before ultimately dying (Fig. 2a; Fig. S3). Bleaching initiated around 4 °C-weeks, peaked around 9 °C-weeks, and then declined sharply thereafter as bleached colonies died (Fig. 2c). Mortality onset was evident from approximately 5 °C-weeks (DHW), with 50% of colonies dead by 10 °C-weeks and almost all dead by 15 °C-weeks (Fig. 2b). Larger colonies (yellow, > 15cm) tended to survive to higher cumulative heat stress levels compared to smaller colonies (purple, < 5cm).

**Fig. 2.**
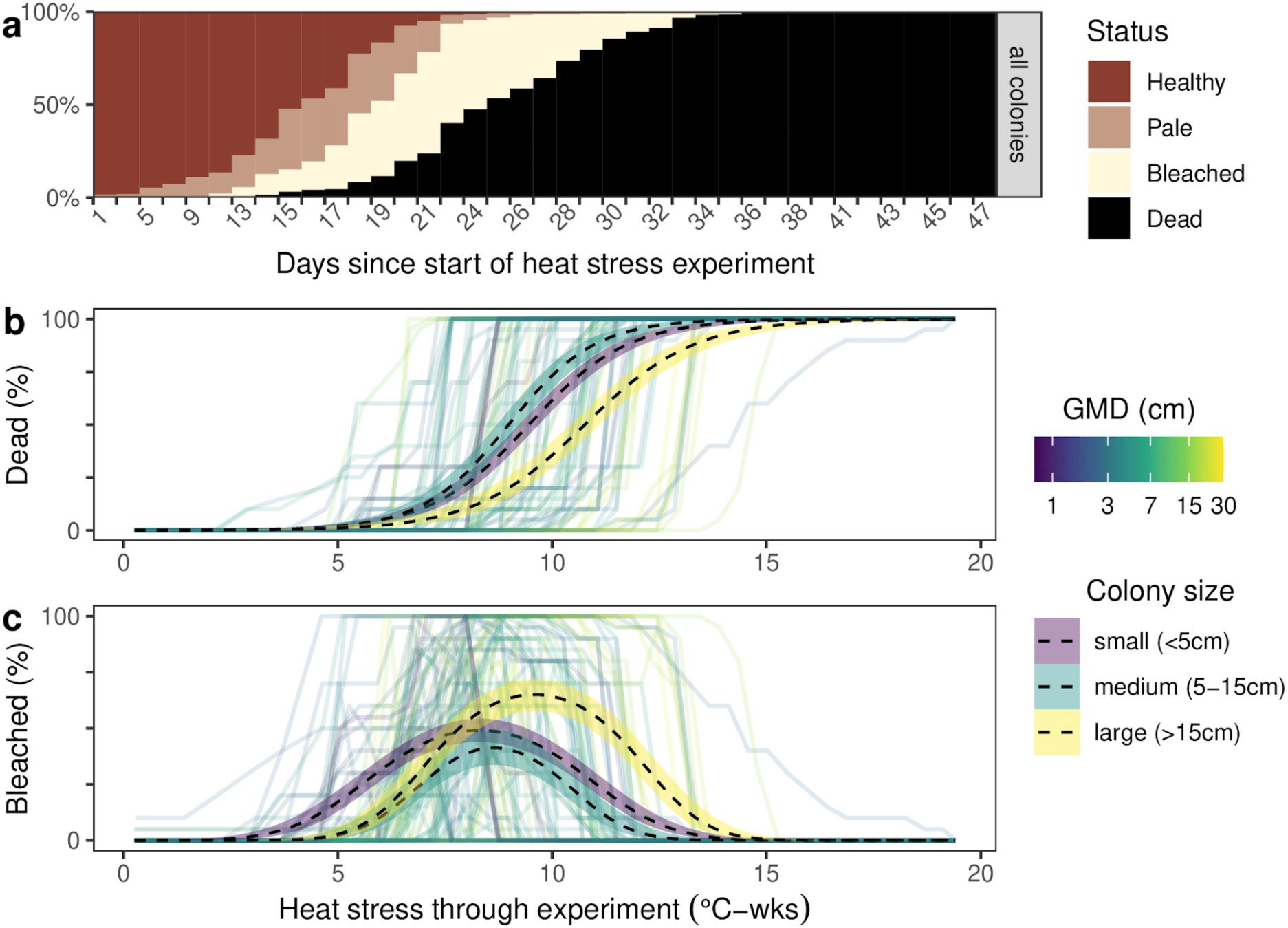
Coral phenotypic status, bleaching, and mortality responses of *A.* aff*. digitifera* across the heat stress experiment. **(a)** Proportion of colonies in each health state (healthy, pale, bleached, dead) over the course of the experiment (days since start of heat stress). **(b)** Individual colony mortality percentage and size-class smoothed trends (dashed lines ± 95% CI) as a function of cumulative heat stress (degree heating weeks (DHW), °C-wks). **(c)** Individual colony bleaching percentage and size-class smoothed trends as a function of cumulative heat stress (degree heating weeks (DHW), °C-wks). In (b) and (c), line color represents colony geometric mean diameter (GMD, cm) on a continuous scale, purple = small (<5 cm), light blue = medium (5–15 cm), and yellow = large (>15 cm) size classes.

### 3.3 Size-dependent survival trajectories

Colony mortality probability increased strongly with cumulative heat stress and was significantly affected by colony size. As expected, there was a significant positive effect of DHW on mortality (binomial GLMM, β = 0.97 ± 0.06, z = 15.89, p < 0.001; Table S2). Colony size (GMD) also had a significant negative effect on mortality (β = −0.18 ± 0.07, z = −2.66, p = 0.008), such that larger colonies were less likely to die at any given level of heat stress. The interaction between DHW and GMD was associated with a higher level of uncertainty (β = 0.010 ± 0.006, z = 1.70, p = 0.090).

Model-predicted mortality trajectories illustrated a clear rise in heatwave tolerance with increasing colony size (Fig. 3a). Small colonies (∼2 cm) reached 50% mortality at approximately 8.4 °C-weeks DHW, compared to 9.6 °C-weeks for medium colonies (∼15 cm) and 10.6 °C-weeks for large colonies (∼29 cm), representing a difference of over 2 °C-weeks across the observed size range (Fig. 3a). Colony size had the most varied influence on survival outcomes at a heat stress of 10 °C-weeks (Fig. 3b, c), where predicted mortality for small colonies (∼2 cm) reached ∼ 60–80%, compared to only 0-10% for the largest colonies (∼29 cm). Below this range, absolute mortality risk was low across all size classes, while above it, even large colonies converged toward near-certain mortality.

**Fig. 3.**
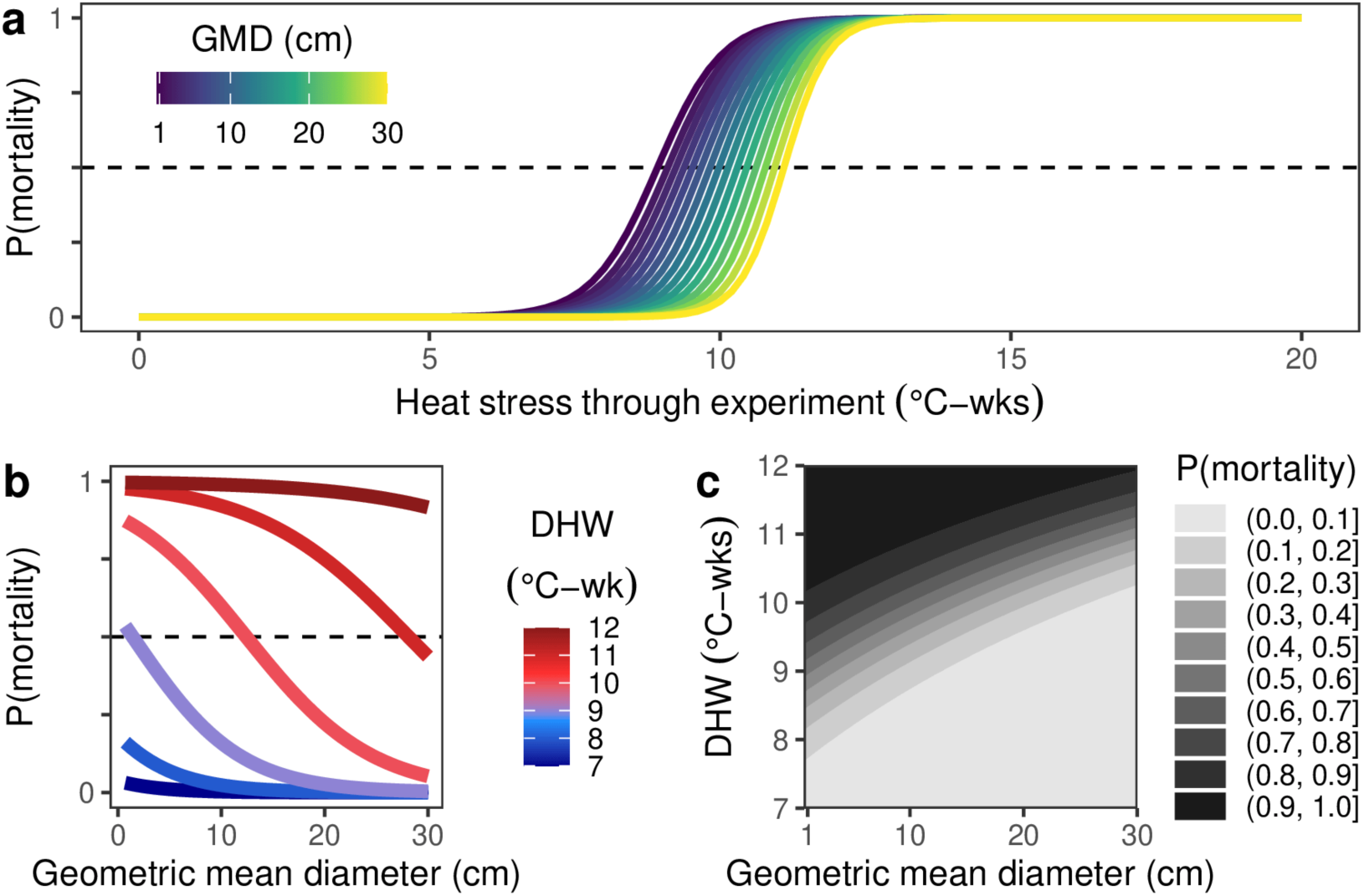
Size-dependent mortality trajectories predicted from a binomial GLMM. **(a)** Predicted probability of mortality as a function of cumulative heat stress (DHW, °C-weeks) across a range of colony sizes (GMD 1–30 cm). **(b)** Predicted probability of mortality as a function of colony size (GMD, cm) at discrete DHW levels (7–12 °C-weeks). **(c)** Filled contour plot of predicted mortality probability across the joint DHW–GMD space (7–12 °C-weeks × 1–30 cm GMD).

### 3.4 Colony size thresholds for bleaching and mortality

The lethal dosage of heat required to elicit 50% chance of mortality (LD50) for a colony was ∼10 °C-weeks on average, but was also highly variable, ranging from approximately 6 to 15 °C-weeks, with most colonies falling between 8 and 12 °C-weeks (Fig. 4a). Segmented regression revealed a significant breakpoint in the relationship between LD50 and colony GMD at 13.4 cm ± 4.4 SE (Fig. 4b). Below this threshold, LD50 remained relatively stable (i.e., 9.5–10 °C-weeks) regardless of colony size, while above it, LD50 increased with colony diameter.

**Fig. 4.**
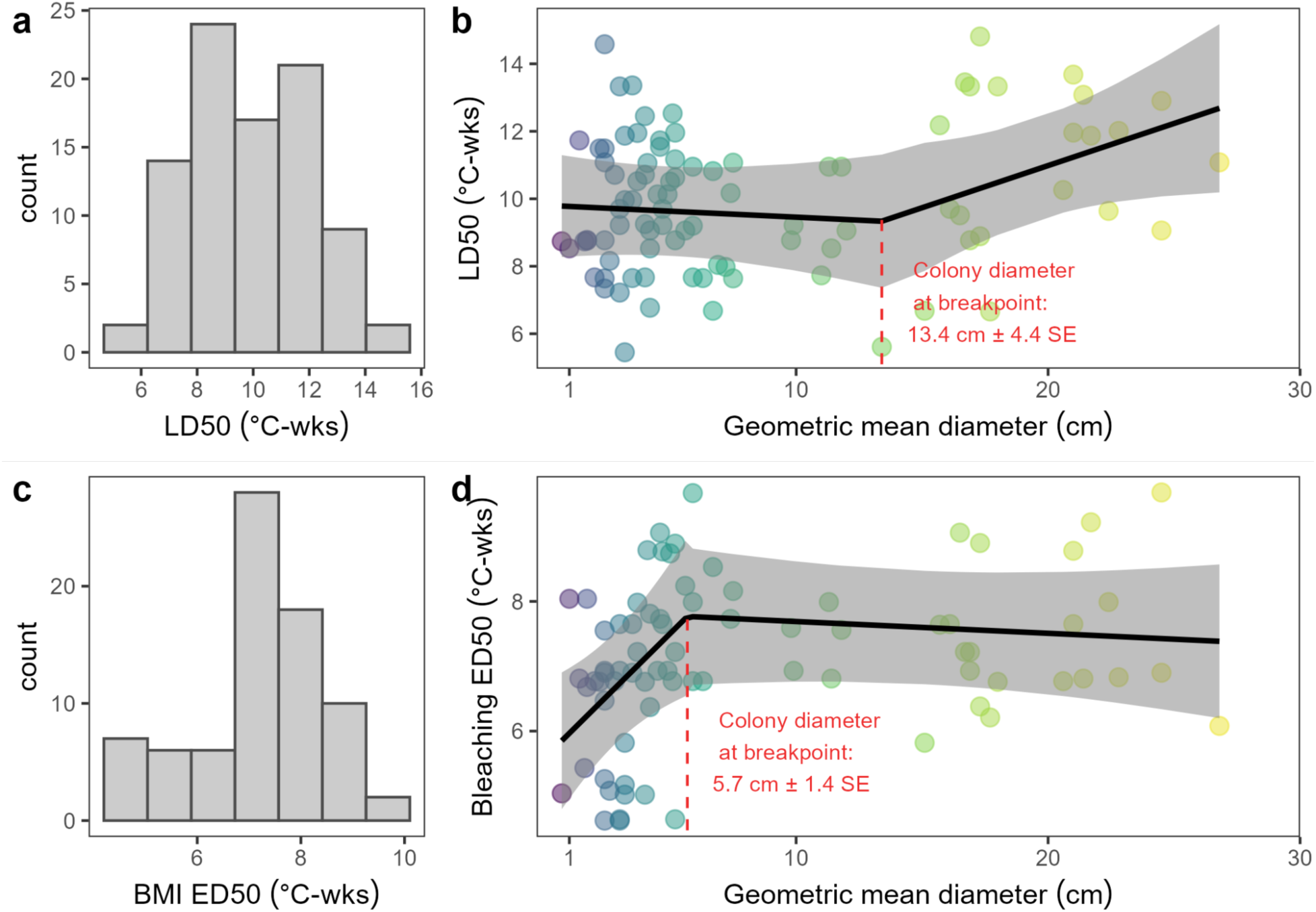
Segmented regression of thermal tolerance endpoints against coral colony size. Histograms show the frequency distributions of LD50 (a) and Bleaching ED50 (c) across colonies. Scatter plots show LD50 (N=87) (b) and Bleaching ED50 (N=77) (d) as a function of colony geometric mean diameter (GMD) on a continuous scale from small (purple) to large (yellow). The solid black line represents the fitted segmented regression model, and the grey shaded ribbon indicates the 95% confidence interval. The red dashed vertical line marks the estimated breakpoint in the relationship between colony size and thermal tolerance, with the corresponding colony diameter and standard error annotated in red.

**Fig. 5.**
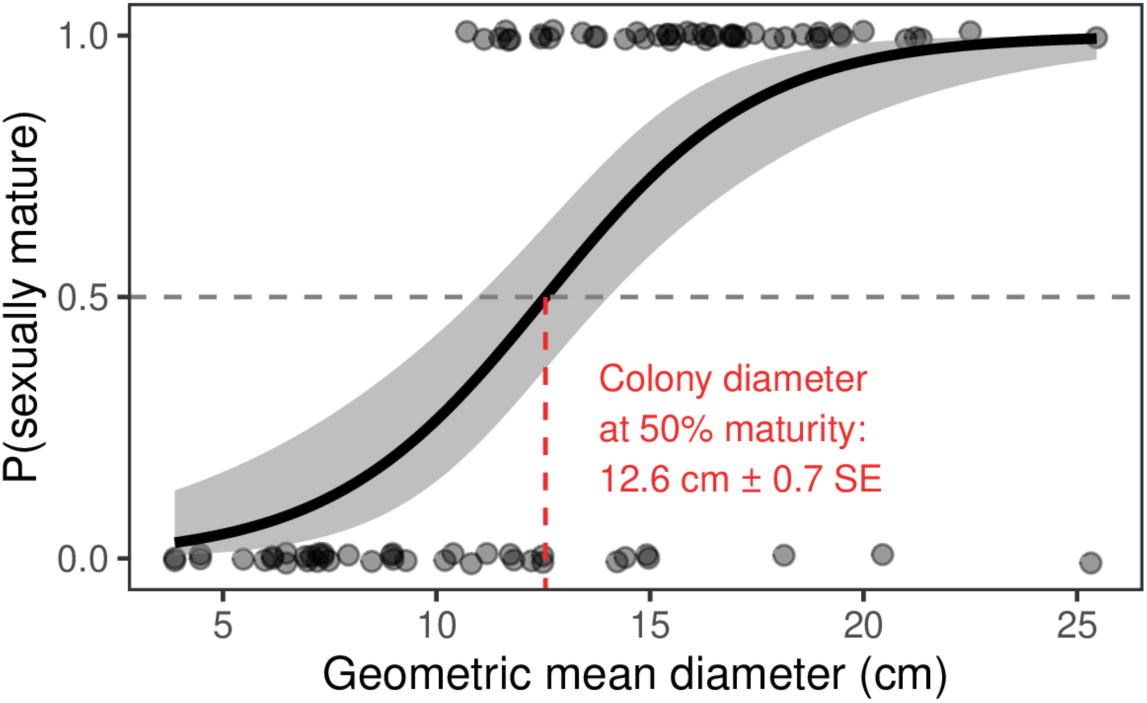
Size threshold for sexual maturity in *A.* aff*. digitifera*. The probability of sexual maturity, P (sexually mature), is plotted as a function of colony geometric mean diameter (GMD). Individual colonies are shown as jittered points (0 = immature, 1 = mature). The solid black line represents the fitted logistic regression model, and the grey shaded ribbon indicates the 95% confidence interval. The horizontal grey dashed line marks the 0.5 probability threshold, and the red dashed vertical line indicates the estimated colony diameter at which 50% of the population reached maturity.

The effective dosage of heat stress required to elicit 50% bleaching (ED50) was more tightly distributed, with most colonies bleaching at 7–9 °C-weeks (Fig. 4c). The segmented model identified a size-based breakpoint for bleaching at a smaller colony size of 5.7 cm ± 1.4 SE GMD (Fig. 4d). Corals below this size exhibited lower tolerance, while those larger showed a higher and stable ED50. Of the 89 colonies observed, 12 were excluded from the bleaching ED50 calculation because they exhibited partial mortality without prior bleaching, leaving 77 colonies for analysis. To incorporate all 89 colonies, the Bleaching Mortality Index (BMI) was also applied, which similarly identified a breakpoint at 4.6 cm ± 1.3 SE GMD (Fig. S4).

### 3.5 Threshold for sexual maturity

Reproductive monitoring of colonies in the field identified that those smaller than 10.7 cm GMD were exclusively immature (Fig. 4). The logistic regression model estimated that 50% of the Mascherchur Reef *A.* aff*. digitifera* population reaches sexual maturity at a colony diameter of 12.6 cm ± 0.7 SE.

## 4. Discussion

In wild coral populations, smaller corals often survive heatwaves better than adults (Burn et al., 2023; Mumby, 1999). However, exposure of corals of diverse sizes to a marine heatwave under controlled light and flow conditions revealed the opposite pattern, with small juveniles showing a greater susceptibility to heat stress than bigger mature colonies. These findings highlight that coral survival in marine heatwave conditions depends on a complex interplay between size-dependent shifts in thermal physiology and localized environmental buffering provided by microhabitats. Our results show that the bleaching response of *A.* aff*. digitifera* started at 4 °C-weeks and peaked at 9 °C-weeks, while the onset of mortality started at 5°C-weeks and peaked at 10 °C-weeks. These thermal thresholds align with values recorded across the Great Barrier Reef for multiple species (Skirving et al., 2020) or previous studies on adults *A.* aff*. digitifera* from the same reef conducted on coral fragments rather than whole colonies (Humanes et al., 2022). Notably, we detected a significant negative effect of size on mortality risk, with smaller colonies (≤10 cm) dying at 10 °C-weeks, whereas sexually mature colonies (≥13.4 cm) maintained a low onset of mortality. Multiple, non-mutually exclusive mechanisms may underlie this pattern, including intrinsic physiological drivers, the protective effect of microhabitats in the reef, or the combination of both factors.

### 4.1 Physiological drivers of size-dependent thermal tolerance

Water flow directly influences the thickness of the coral diffusive boundary layer (DBL), a key determinant of mass transfer rates between coral tissue and the surrounding seawater (Patterson et al., 1991). Studies show that under both laminar and turbulent flow, flat organisms exhibit higher mass transfer rates than thin cylindrical forms (Comeau et al., 2019; Patterson et al., 1991; Shashar et al., 1993, 1996). Faster mass transfer favor the removal of by-products such as superoxide and other oxygen radicals that accumulate under conditions of high temperature and high solar radiation (Lesser, 1997) and has therefore been proposed as a physiological mechanism favoring increased thermal tolerance in small, flat juvenile *Acropora* colonies relative to larger, branching adults (Baird & Atkinson, 1997; Loya et al., 2001; Ng et al., 2019; Shenkar et al., 2005; van Woesik, Irikawa, et al., 2012). To test whether flow-related mass transfer could account for size-dependent thermal tolerance in our system, we used clod cards to confirm that ambient flow conditions did not differ systematically between smaller and larger colonies, ruling out gross differences in water motion as a confounding factor. Yet, our results show that smaller colonies exhibited an earlier onset of both bleaching and mortality than larger colonies when exposed to comparable levels of light and flow, with the probability of dying at 10 °C-weeks being markedly higher in smaller sizes. These findings therefore suggest that differences in mass transfer alone cannot explain the size-dependent thermal responses observed here.

The existence of distinct size thresholds for bleaching (GMD = 5.7 cm) and mortality (GMD = 13.4 cm) suggests that these processes are governed by different physiological mechanisms. Bleaching likely reflects a lower photo-physiological tolerance threshold in colonies smaller than 5.7 cm GMD, whereas the marked reduction in mortality above 13.4 cm GMD indicates that survival during prolonged heat stress depends more strongly on colony energetic balance, particularly the availability of energy reserves before bleaching (Anthony et al., 2009). Notably, this mortality threshold closely matches the size at sexual maturity of the wild *A.* aff*. digitifera* population of this study, suggesting that key ontogenetic transitions coincide with changes in thermal resilience and tolerance. Juvenile colonies primarily allocate resources towards somatic growth and tissue expansion (van der Meer, 2006). Structural lipids, such as sterols and membrane phospholipids enriched in polyunsaturated fatty acids (PUFAs), play a central role in cell membrane synthesis, cellular development, and somatic growth (Sikorskaya et al., 2025). As colonies mature, lipid metabolism shifts toward accumulation of neutral storage lipids, particularly triacylglycerols and wax esters, which fuel gametogenesis (Arai et al., 1993; Harrison, 2011; Leuzinger et al., 2012) while providing a readily mobilized energy reserve during periods of stress (Anthony et al., 2009; Harii et al., 2007; Oku et al., 2002). Unlike sterols, which cannot be catabolized because of their chemically stable ring structures, or PUFAs, which require additional enzymatic processing before β-oxidation, triacylglycerols and wax esters can be rapidly hydrolyzed and metabolized to sustain respiration under carbon limitation during heat stress (Ermolenko & Sikorskaya, 2021; Imbs, 2013; Rodrigues & Grottoli, 2007). Consequently, reproductively mature colonies are likely to possess both larger and more metabolically accessible energy reserves than juveniles.

These ontogenetic differences in lipid metabolism are consistent with previous studies showing that adult corals maintain larger tissue energy reserves and increasingly rely on heterotrophic carbon acquisition to sustain metabolism during bleaching (Grottoli et al., 2006; Meunier et al., 2019, 2022; Rodrigues & Grottoli, 2007). Although thermal stress induces substantial remodeling of membrane lipid composition, bulk storage lipids are generally depleted only during prolonged bleaching (Sikorskaya et al., 2025). Consequently, the quantity and composition of lipid reserves accumulated before bleaching onset are likely to determine how long colonies can maintain basal metabolism after the loss of symbiont-derived carbon (Anthony et al., 2009). Together, these mechanisms provide a plausible explanation for why sexually mature colonies in our experiment survived substantially longer than juveniles.

Another intrinsic mechanism that could contribute to size-dependent thermal tolerance is ecological memory mediated through epigenetic processes. Environmental stress can induce epigenetic modifications that alter gene expression without changing DNA sequences, potentially enhancing acclimatization to future heat stress (Putnam et al., 2020; Putnam & Gates, 2015). Larger colonies, having experienced longer and more variable environmental exposure, may therefore possess a stronger ecological memory and enhanced thermal resilience (Hackerott et al., 2021). In *A.* aff*. digitifera*, colonies of approximately 40 cm diameter from the same reef have been estimated to be approximately 12 years old (Lachs et al., 2025), while reproductive maturity is typically reached after 3 to 4 years (Lachs et al., 2026; Villanueva et al., 2012). However, because major bleaching events in Palau were last recorded in 1998 and 2010 (Bruno et al., 2001; van Woesik, Houk, et al., 2012), colonies selected for this experiment were unlikely to have experienced prolonged extreme marine heatwaves during most of their lifespan. Consequently, epigenetic priming associated with repeated thermal stress exposure is unlikely to explain the observed patterns in this case.

### 4.2 Microenvironmental conditions determining size-dependent thermal tolerance

While our experimental design indicates that intrinsic physiological changes accompanying size contribute to the increase in thermal tolerance with colony size, these mechanisms are difficult to isolate in wild populations because juvenile and adult corals experience markedly different microhabitats. Consequently, apparent differences in thermal tolerance in the field may reflect not only physiological changes associated with colony size but also differences in the microenvironmental conditions to which colonies are exposed.

Coral larvae preferentially settle within micro-refugia in response to both physical (e.g., light availability and hydrodynamic conditions) and biological factors (e.g., their initial aposymbiotic state, protection from corallivory, and reduced benthic competition) (Arnold et al., 2010; Doropoulos et al., 2015; Randall et al., 2021). As a result, newly settled corals are typically found in cryptic or shaded habitats, where exposure to ultraviolet radiation (UVR) and photosynthetically active radiation (PAR) is substantially lower than that experienced by exposed adult colonies. Because elevated irradiance acts synergistically with thermal stress to induce bleaching (Brown, 1997), these early-life habitats are likely to reduce the cumulative light stress experienced by juvenile colonies during marine heatwaves (Mumby, 1999). Consequently, juveniles and adults are exposed to fundamentally different thermal and light microenvironments, potentially biasing direct comparisons of bleaching susceptibility from field observations.

This environmental mismatch may help reconcile the apparent discrepancy between our experimental results, which indicate lower intrinsic thermal tolerance in juvenile colonies, and numerous field studies reporting greater bleaching resistance in early life stages (Álvarez-Noriega et al., 2018; Bena & Van Woesik, 2004; Brandt, 2009; Mumby, 1999; Speare et al., 2022). Rather than representing contradictory findings, our results highlight a possible compensatory interaction between the two factors: colony size has a negative effect on thermal tolerance, while microrefugia may confer a strong positive benefit. In the field, where both factors operate simultaneously, their effects appear to combine additively, with the protective benefit of microrefugia exceeding the physiological vulnerability associated with smaller colony size.

### 4.3 Ecological and conservation implications

Because intrinsic thermal tolerance and effective thermal exposure are jointly determined by colony size and habitat, changes in reef structural complexity under climate change may disproportionately affect juvenile survival. The survival advantage conferred to larger, sexually mature colonies means that reef populations retaining a sufficient proportion of adult individuals are more likely to maintain reproductive output following bleaching events, providing the larval supply necessary for subsequent recruitment and recovery, once the latent effects of bleaching on coral physiology have been recovered (Briggs et al., 2024; Johnston et al., 2020). Our findings suggest that juvenile corals possess a reduced physiological capacity to tolerate marine heatwaves, although this may not necessarily translate into higher mortality when reef structural complexity provides sufficient environmental buffering or across other coral taxa. Indeed, shaded microhabitats may buffer juveniles from the synergistic effects of high irradiance and thermal stress, suggesting that incorporating substrate heterogeneity or habitat complexity into size-dependent simulation models of coral populations could improve the accuracy of future projections under climate change.

Reproductively mature colonies contribute to reef structural complexity and provide important habitat that supports diverse reef-associated communities and ecosystem functions (Done et al., 1996). Our study indicates that sexually mature colonies exhibit increased thermal tolerance, a finding with direct implications for climate-smart management strategies and coral reef conservation. Size structure could therefore serve as a practical demographic indicator of reef resilience, helping managers identify populations with greater recovery potential following marine heatwaves. In conservation contexts, these findings highlight the importance of conserving adult broodstock, minimizing disturbances that disproportionately remove large colonies, and implementing mitigation measures during marine heatwaves. Lastly, incorporating colony size distributions into monitoring and restoration planning may improve predictions of reef persistence under continued ocean warming.

### 4.4 Study limitations

The objective of this study was to quantify the influence of coral colony size on heatwave-driven bleaching and mortality. While adult colonies of *A.* aff*. digitifera* possess distinct morphological traits that facilitate species-level identification (e.g., colony morphology, radial and axial corallite shape, and axial corallite coloration), the identification of juvenile colonies remains challenging because juveniles typically exhibit flatter and less differentiated morphologies. To reduce taxonomic uncertainty, colony selection was guided using reference photographs of juvenile *A.* aff*. digitifera* previously reared in both in situ and ex situ nurseries (see Fig. S5). Despite the authors having worked with multiple life stages of this species at the study site for more than eight years, generating extensive observational records of phenotypic variation throughout ontogeny, we can not exclude the possibility that some juveniles were misidentified. Molecular genetic analyses would be required to confirm species identity. However, tissue sampling was intentionally avoided, particularly for small colonies, to minimize any potential effects on bleaching and mortality responses during the experiment.

Additionally, although *A.* aff. *digitifera* is a widely distributed and common species throughout the Indo-Pacific, we must acknowledge that our study was conducted on a single corymbose species. We believe that our findings are going to be relevant to other broadcast-spawning corals (∼80% of the known coral species), which share similar modes of horizontal symbiont acquisition. However, it is not clear if our conclusions can apply to species with vertical transmission of algal symbionts, as their symbiont communities are established before release and may confer different thermal physiology across life stages. However, while several studies report elevated thermal tolerance in juveniles of *Acropora*, comparable work has documented increased bleaching susceptibility in juveniles of *Pocillopora* and *Montipora* relative to conspecific adults, both of which rely on vertical symbiont transmission (Álvarez-Noriega et al., 2018; Burn et al., 2023). Further studies on vertically transmitted species are needed to resolve this current research gap.

Together, our findings demonstrate that intrinsic thermal tolerance increases with colony size in *A.* aff. *digitifera*, with a notable increase at the onset of sexual maturity. These results suggest that the vulnerability of juvenile corals on natural reefs ultimately reflects an interaction between physiology and microhabitat: colony size negatively affects intrinsic thermal tolerance in juveniles, while microrefugia buffer this vulnerability and delay bleaching onset. Understanding both components will be essential for accurately predicting coral population responses to increasingly frequent marine heatwaves.

## Acknowledgments

Authors would like to thank Arius Merep, Geory Mereb, and all the rest of PICRC staff who helped with the logistics during the experiment. This study was funded by Coral Research and Development Accelerator Platform (CORDAP), and corals were collected under permit #26-03 issued by Palauan national authorities.

